# Characterization of a selective, iron-chelating antifungal compound that disrupts fungal metabolism and synergizes with fluconazole

**DOI:** 10.1101/2023.08.09.552590

**Authors:** Jeanne Corrales, Lucía Ramos-Alonso, Javier González-Sabín, Nicolás Ríos-Lombardía, Nuria Trevijano-Contador, Henriette Engen Berg, Frøydis Sved Skottvoll, Francisco Moris, Óscar Zaragoza, Pierre Chymkowitch, Ignacio Garcia, Jorrit M. Enserink

## Abstract

Fungal infections are a growing global health concern due to the limited number of available antifungal therapies as well as the emergence of fungi that are resistant to first-line antimicrobials, particularly azoles and echinocandins. Development of novel, selective antifungal therapies is challenging due to similarities between fungal and mammalian cells. An attractive source of potential antifungal treatments is provided by ecological niches co-inhabited by bacteria, fungi and multicellular organisms, where complex relationships between multiple organisms has resulted in evolvement of a wide variety of selective antimicrobials. Here, we characterized several analogs of the one such natural compound, Collismycin A. We show that NR-6226C has antifungal activity against several pathogenic *Candida* species, including *C. albicans* and *C. glabrata*, whereas it only has little toxicity against mammalian cells. Mechanistically, NR-6226C selectively chelates iron, which is a limiting factor for pathogenic fungi during infection. As a result, NR-6226C treatment causes severe mitochondrial dysfunction, leading to formation of reactive oxygen species, metabolic reprogramming and a severe reduction in ATP levels. Using an *in vivo* model for fungal infections, we show that NR-6226C significantly increases survival of *Candida*-infected *Galleria mellonella* larvae. Finally, our data indicate that NR-6226C synergizes strongly with fluconazole in inhibition of *C. albicans*. Taken together, NR-6226C is a promising antifungal compound that acts by chelating iron and disrupting mitochondrial functions.

**Importance statement:** Drug-resistant fungal infections are an emerging global threat, and pan-resistance to current antifungal therapies is an increasing problem. Clearly, there is a need for new antifungal drugs. In this study, we characterized a novel antifungal agent, the Collismycin analog NR-6226C. NR-6226C has a favorable toxicity profile for human cells, which is essential for further clinical development. We unraveled the mechanism of action of NR-6226C and found that it disrupts iron homeostasis and thereby depletes fungal cells of energy. Importantly, NR-6226C strongly potentiates the antifungal activity of fluconazole, thereby providing inroads for combination therapy that may reduce or prevent azole resistance. Thus, NR-6226C is a promising compound for further development into antifungal treatment.

## Introduction

Fungal infections range from relatively benign infections of the skin and mucosal tissues to life-threatening invasive infections, affecting more than a billion people worldwide and killing over 1.5 million people annually^1^. Infections with *Candida* species, such as *C. albicans* and *C. glabrata*, are among the most common human fungal infections^1^. Among the main antifungals that are used in the clinic today are azoles, echinocandins, and polyenes^2^. Azoles block cell membrane synthesis by inhibiting the ergosterol biosynthesis enzyme, lanosterol 14-α-demethylase (encoded by *ERG11* in *Candida*), whereas echinocandins disrupt cell wall synthesis via noncompetitive inhibition of the (1,3)-β-d-glucan synthase enzyme encoded by *FKS* genes. However, long-term exposure to antifungals has resulted in a steadily increasing prevalence of antifungal resistance, which is a major global health concern^3^. Therefore, there exists a need for new antifungal drugs for treatment of antifungal-resistant *Candida* and other pathogenic fungal species.

During evolution, dynamic interactions that occur in ecological niches co-inhabited by bacteria and fungi have resulted in formation of a wide range of antimicrobial compounds that are a rich source of potential antifungal treatments^4^. One example of a complex multi-organism interaction occurs in colonies of leafcutter ants, which form a mutually beneficial symbiotic relationship with a specialized fungus and with various antimicrobials-producing bacteria, including *Streptomyces sp*. The ants provide freshly cut leaves to the fungus, resulting in a fungal garden. The fungi produce specialized fungal structures that serve as a key food source for the ant larvae^5^. The bacteria, which grow on the cuticle of the ants, synthesize antifungal compounds such as candicidin or nystatin that protect the fungal garden from invasion by parasitic fungi, which can ruin the fungal garden and destroy the colony^6, 7^. Another such compound is Collismycin A (ColA; Fig. 1A), which is produced by *Streptomyces* and which has antimicrobial activity against several fungi, including *C. albicans*^8^. However, ColA also possesses substantial cytotoxic activities against mammalian cells^9^, rendering it less useful as antifungal therapy.

**Figure 1.**
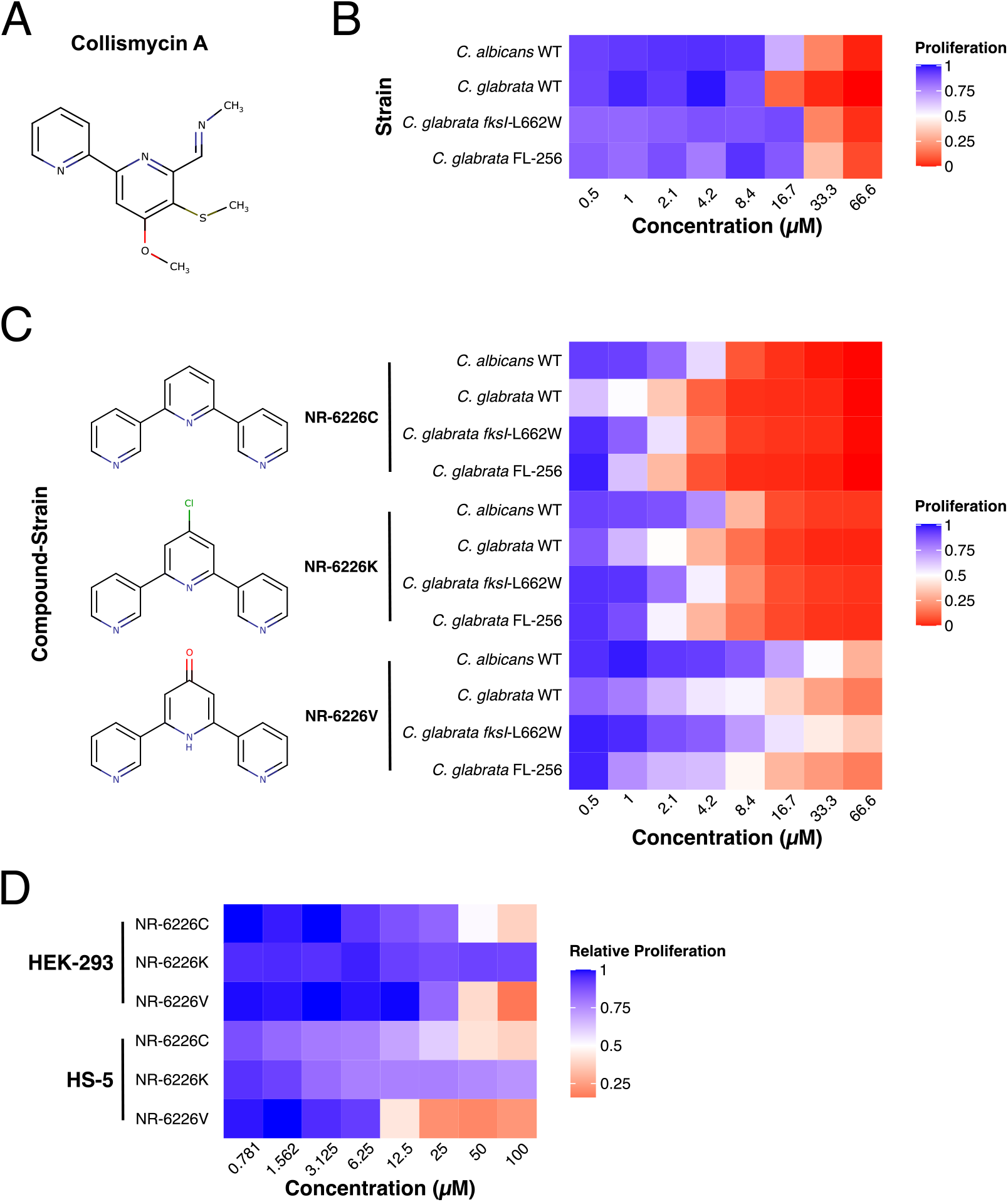
Identification of the ColA analog NR-6226C as an antifungal agent with a potential therapeutic window. ***A***, ColA compound structure. ***B*,** Heat map of relative proliferation of several WT and echinocandin- and azole-resistant *Candida* strains. The indicated strains were incubated in SD media with increasing concentrations of ColA for 24h, after which proliferation was analyzed by measuring the OD_600_. Data were normalized to DMSO-treated control samples. ***C,*** Heat map of *Candida* proliferation in the presence of the ColA analogs NR-6226C, NR-6226K, and NR-6226V. The indicated fungal strains were incubated in the presence of increasing compound concentrations in YNB-iron media for 24h, after which proliferation was measured using OD_600_. ***D,*** Human HEK-293 and HS-5 cells are considerably less sensitive to ColA analogs than *Candida spp*. Cells were incubated with increasing concentrations of the indicated compounds for 24 hrs, after which relative cell viability was assayed using CellTiter-Glo. Data were normalized to DMSO controls.

In this study, we characterized the antifungal activity and overall toxicity of several ColA analogs and identified a compound with antifungal activity against clinical isolates of azole- and echinocandin-resistant *Candida* species, but with reduced toxicity against mammalian cells. Our results suggest that this ColA analog is a promising candidate for development into treatment of infections caused by wild-type and drug-resistant *Candida* species.

## Results

### Identification of Collismycin A analogs with increased antifungal activity

To investigate the potential use of novel ColA analogs as selective antifungal agents, we generated a compendium of 26 collismycin-related compounds (Suppl. Fig. S1A; see Methods for details). To determine selectivity against pathogenic fungi, we screened these compounds against a panel of *Candida* strains and two human cell lines. We used *Candida albicans* (CCUG32723), *Candida glabrata* (ATCC15545), as well as an echinocandin-resistant *C. glabrata* strain (*fksI-L662W*; JEY12725) and a fluconazole-resistant *C. glabrata* strain (FL-256; JEY12726), both of which were isolated at Oslo University Hospital (Suppl. Table S1). First, we studied the proliferation of these fungi on increasing concentrations of the starting compound, ColA. ColA reduced the proliferation of all pathogenic yeasts tested (Fig. 1B; Suppl. Fig. S2 and S3A). We then tested the other analogs and found that the antifungal activity was lost for most of the compounds (Suppl. Fig. S2). However, three compounds, NR-6226C, NR-6226K, and NR-6226V inhibited the growth of *Candida spp* more potently than ColA (Fig. 1C, Suppl. Fig. S3A and B). NR-6226C appeared to be the most efficacious, yielding an EC_50_ at least 12 times smaller than ColA-treated strains, whereas treatment with either NR-6226K or NR-6226V resulted in an inhibitory effect that was approximately 6 times greater than ColA (Suppl. Fig. S3B). Importantly, the two clinically isolated antimicrobial-resistant *C. glabrata* strains were also sensitive to these compounds (Fig. 1C and Suppl. Fig. S3B). These results encouraged us to further explore the potential of these drugs for further development into antifungal therapy.

For any novel antifungal compound to have a clinical application, it is essential that there exists a therapeutic index, i.e. fungi should be more sensitive to the compound than mammalian cells. Therefore, we tested the sensitivity of HS-5 and HEK-293 fibroblast cell lines to NR-6226C, NR-6226V and NR-6226K. We found that NR-6226V strongly attenuated cell proliferation of HS-5 cells (10.49 ± 1.42 µM), and to a somewhat lesser extent also that of HEK-293 cells, limiting its usefulness as a potential antifungal treatment (Fig. 1D and Suppl. Fig. S3C). In contrast, EC_50_ values of NR-6226C were 37.05 ± 6.94 µM and 28.68 ± 8.32 µM in HEK-293 and HS-5 cell lines, respectively (Fig. 1D and Suppl. Fig. S3C), and although we noticed morphological changes at very high concentrations of NR-6226C (100 µM; Suppl. Fig. S4), these data suggest the existence of a potential therapeutic window. Treatment with NR-6226K appeared to have no discernible effects on the proliferation of either cell lines; however, closer inspection revealed precipitation of the compound (Suppl. Fig. S4), rendering NR-6226K less useful in physiological settings. Together, these results show that the ColA analogs NR-6226C and NR-6226K have antifungal activity exceeding that of ColA. For the remainder of the study, we decided to focus on NR-6226C due to its superior activity compared to ColA, its better solubility, its favorable toxicity profile for human cells and its effectiveness against antifungal-resistant *C. glabrata* strains.

### ColA analogue NR-6226C is a Fe^2+^ chelator

ColA has been previously shown to have iron-chelating properties through the formation of a complex containing two ColA molecules bound to a single atom of either Fe^2+^ or Fe^3+^, and the three nitrogen atoms in ColA have been proposed to facilitate binding of the iron atom to form a tridentate chelator^9, 10, 11^.

To test whether iron chelation is important for the inhibitory effect of ColA on fungal growth, we tested the antifungal activity of Collismycin H (ColH), which is a ColA derivative that contains a hydroxymethyl instead of a methyl-imine moiety and therefore cannot chelate Fe^2+/3+^ (Fig. 2A, *right*). As shown in Figure 2A, ColH lacked antifungal activity even at very high concentrations, showing that the three nitrogen atoms in ColA are essential for antifungal activity. While not conclusive by themselves, these data do support the idea that ColA antifungal activity and its derivatives is mediated through iron chelation. Given that some metal chelators can be promiscuous and bind different metals with different affinities^12^, we tested which metals can be bound by NR-6226C using high-performance liquid chromatography and mass spectrometry. These nano liquid chromatography experiments showed that similar to ColA, NR-6226C forms a 2:1 compound-iron complex, where it preferentially binds Fe^2+^ in a concentration-dependent manner, and to a much lesser extent also Fe^3+^ (Fig. 2B and Suppl. Fig. S5).

**Figure 2.**
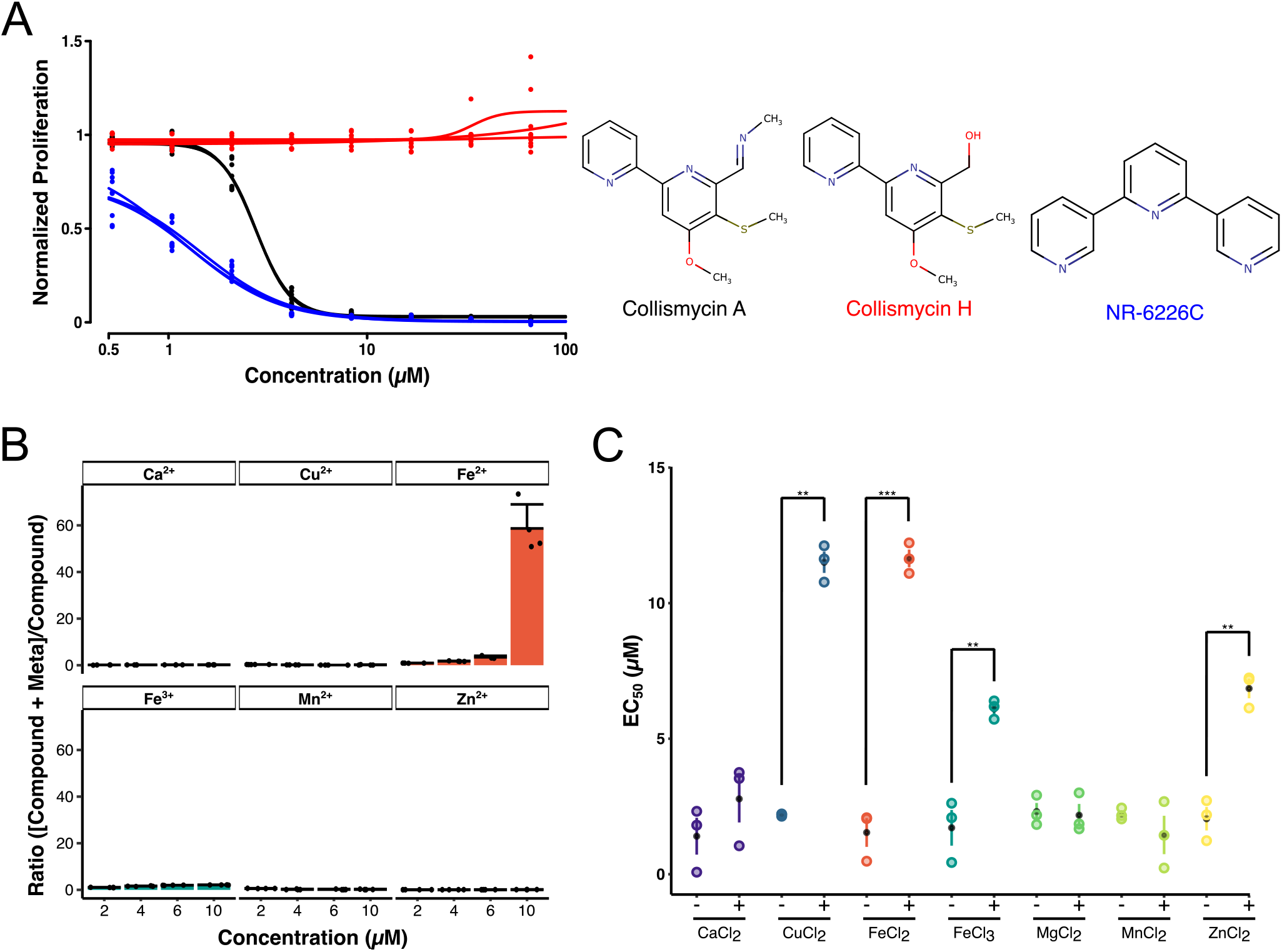
NR-6226C is an iron chelator that inhibits *Candida* proliferation by sequestering iron. ***A***, ColH, which does not bind iron, does not inhibit growth of *C. glabrata.* Fungal cells were treated with ColA, ColH and NR-6226C in SD media for 24h, after which proliferation was measured using OD_600_. Data were normalized to DMSO control samples. ***B,*** NR-6226C selectively binds Fe^2+^ *in vitro*. The interaction of NR-6226C with increasing concentrations of the indicated metal ions was determined as described in Methods. *n=3*. ***C,*** Exogenous Fe^2+^ rescues the antiproliferative effect of NR-6226C. Wild-type *C. glabrata* was incubated for 24h with NR-6226C in presence of 5 µM CaCl_2_, CuCl_2_, FeCl_2_, FeCl_3_, MgCl_2_, MnCl_2_, or ZnCl_2_, respectively. Proliferation was measured using OD_600_ and values were normalized to DMSO controls. Resulting EC_50_ values were calculated and statistical significance was analyzed using unpaired two-sample t-tests in R, \*\*\**P*≤0.001; \*\**P*≤0.01; *n=3*.

Next, we determined whether chelation of metal ions is important for antifungal activity of NR-6226C. We performed growth assays of *C. glabrata* with NR-6226C in absence or presence of 5 µM of various metal salts. As expected, the addition of FeCl_2_ significantly increased the EC_50_ of NR-6226C (Fig. 2C). The same was observed for FeCl_3_, which is likely due to the fact that Fe^3+^ can be converted into Fe^2+^ by ferric reductases. Surprisingly, Cu^2+^ and Zn^2+^, which hardly bind NR-6226C even at high concentrations, also significantly increased the EC_50_ of NR-6226C. The ameliorating effect of Cu^2+^ and Zn^2+^ on inhibition of fungal growth by NR-6226C may be caused by mismetallation, i.e. inactivation of a protein due to binding of a non-cognate metal ^13, 14^, which has previously been shown to induce a compensatory response that promotes uptake of Fe^2+^ ^15^. Overall, these findings indicate that NR-6226C inhibits the proliferation of *Candida* strains mainly through iron chelation, although we cannot exclude the possibility that it acts through alternative mechanisms *in vivo*.

### Treatment with NR-6226C induces an iron starvation response

We wished to gain more insight into the physiological response of fungal pathogens to NR-6226C treatment by studying changes in gene expression programs. To avoid potential secondary effects of long-term drug treatment, we treated *C. glabrata* cells for 1 hour with 10 µM NR-6226C and analyzed changes in mRNA levels by RNA sequencing. For comparison, we also included treatment with 5 µM Dp44mT, which is a known iron chelator^16^. We found that 224 genes were significantly upregulated and 220 genes were significantly downregulated upon NR-6226C treatment (Suppl. Table S2). Interestingly, genes that were significantly upregulated by NR-6226C included genes involved in the response to iron starvation and the oxidative damage response, such as *TRR1*, *HMX1*, and *HBN1*, which encode thioredoxin, heme oxygenase and an oxidoreductase, respectively (Fig. 3A). Other examples of genes induced by NR-6226C were *CAGL0G06798g*, which is a homolog of *Saccharomyces cerevisiae (Sc) LSO1* and known to be induced by iron starvation^17^; CAGL0L11990g, a homolog of *Sc GRX4*, encoding an iron-response regulating enzyme with glutathione-dependent oxidoreductase and glutathione S-transferase activities^18^; and *CAGL0D04708g*, a major copper transporter that is transcriptionally induced at low copper levels^19^. Genes that were downregulated by NR-6226C include genes encoding proteins with iron-sulfur clusters, such as *CAGL0M00374g*, which encodes H2S-NADP oxidoreductase; the succinate dehydrogenase-encoding gene *SDH2* and the aconitase-encoding gene *ACO1*, which are preferentially expressed in the presence of sufficiently high iron levels^20^. Other downregulated genes are *DUG3*, which mediates degradation of glutathione^21^; and the detoxifying metallothionein gene *MT-I*, which is induced by high levels of metal ions^22^. As expected, gene ontology (GO) analysis revealed that these genes function in mitochondrial and enzymatic functionalities, such as lyase activity, oxidoreductase activity, and catalytic activity (Fig. 3B and C). Together, these findings strongly suggest that NR-6226C treatment induces an iron starvation response and possibly also an oxidative damage response.

**Figure 3.**
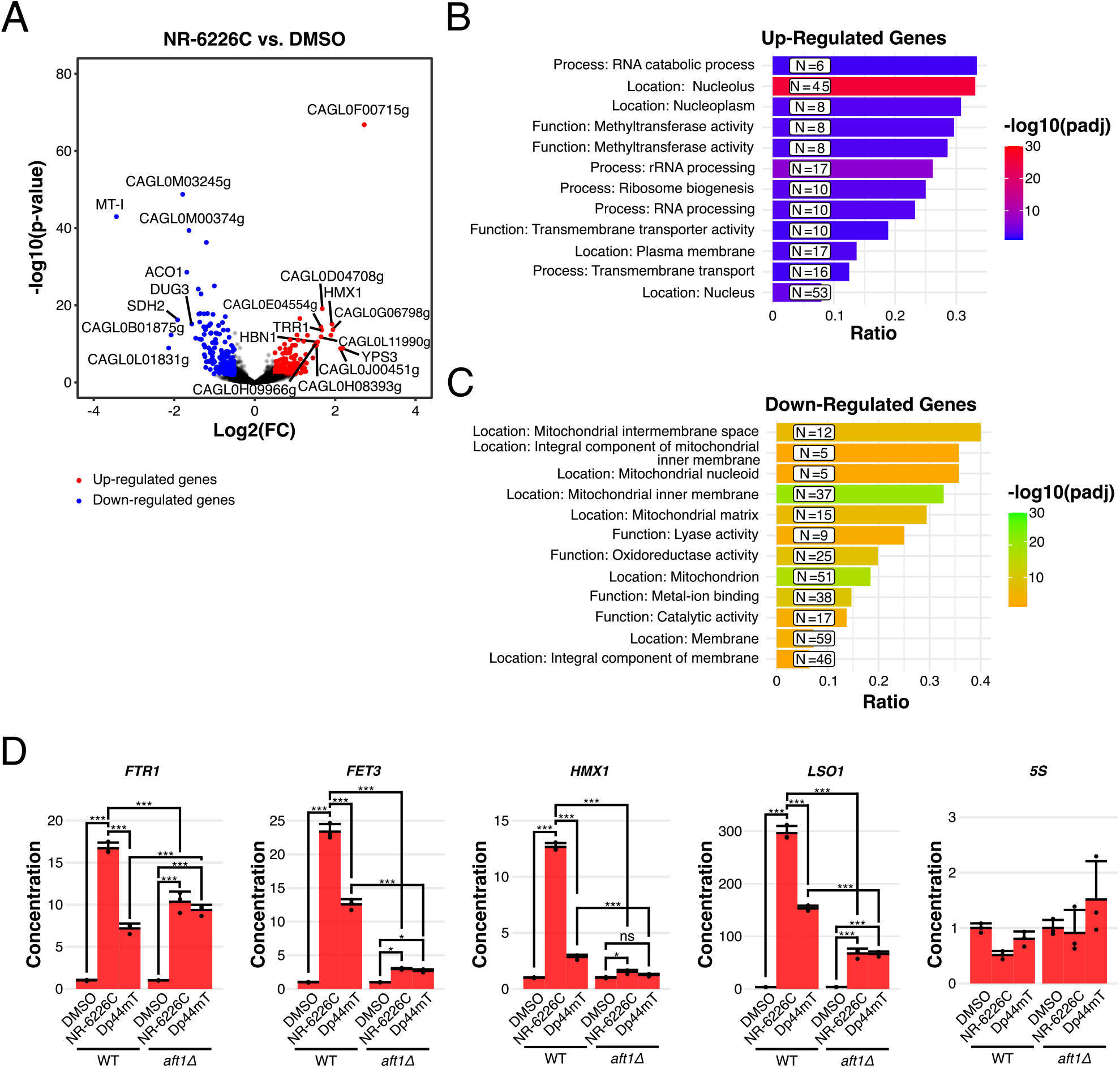
NR-6226C treatment induces an iron starvation response. ***A,*** Wild-type *C. glabrata* cells were incubated with either DMSO or 10 µM NR-6226C for 1h, after which RNA levels were analyzed by RNA-seq. The volcano plot shows transcripts below (blue) or above (red) a 1.4-fold change (log2-fold change <-0.5 or >0.5) and an adjusted p-value <0.01 threshold in NR-6226C-treated cells relative to the DMSO control. ***B***, GO analysis of genes that are either upregulated (top) or downregulated (bottom) by NR-6226C treatment. ***C,*** Validation of a panel of selected genes by RT-qPCR shows that NR-6226C treatment activates the transcription factor Aft1. Wild-type or *aft1*Δ *S. cerevisiae* strains were treated with DMSO, 10 µM NR-6226C, or 10 µM Dp44mT for 24 hrs, after which RNA levels were analyzed by RT-qPCR as described in Methods. cDNA concentrations were normalized to the respective WT and mutant samples treated with DMSO, followed by calculation of the means, sample standard deviations and statistical significance using unpaired two-sample t-tests in base R. \*\*\**P*≤0.001; \*\**P*≤0.01; *ns*, not significant; *n=3*.

To further characterize the transcriptional response of NR-6226C, we compared it to that of the known metal chelator Dp44mT, which chelates both copper and iron^16, 23^. As expected, there was a substantial correlation between the transcriptional responses of these two compounds, showing an overlap of 55% and 58% of genes that were significantly downregulated and upregulated, respectively (Supp. Fig. S6A-C). Genes encoding iron-binding metalloenzymes or enzymes containing iron-sulfur clusters were strongly downregulated by NR-6226C and Dp44mT treatment, such as *SDH2*, *ACO1/2*, *CAGL0E05676g* (*Sc TYW1*), *LYS9*, as well as the iron-sulfur assembly coding gene *CAGL0G03905g* (*Sc ISA1).* These results further support the idea that NR-6226C is a potent iron chelator.

The iron starvation response in *S. cerevisiae* has been reported to involve the transcription factors Aft1 and Aft2^24–26^. Given that *C. glabrata* is very closely related to *S. cerevisiae*^27^, and because it is challenging to create gene deletions in *C. glabrata*, we used *S. cerevisiae* to test whether some of the responses to NR-6226C are mediated by Aft1. For comparison, we included Dp44mT as a control. Interestingly, treatment of wild-type (WT) *Sc* cells with NR-6226C resulted in robust activation of Aft1/2 target genes *FTR1*, *FET3*, *HMX1*, and *LSO1*, while there was no effect of NR-6226C on expression of control 5S RNA (Fig. 3D). Importantly, deletion of the *AFT1* gene attenuated the transcriptional responses upon treatment with NR-6226C (Fig. 3D).

Taken together, we conclude that treatment with NR-6226C induces a strong iron starvation response in *C. glabrata* and *S. cerevisiae*. Given that iron is a limiting factor in microbial pathogenesis^28^, NR-6226C is a promising candidate for treatment of microbial infections.

### Treatment with NR-6226C induces ROS formation

Low levels of iron can result in dysregulation of iron-dependent processes, including oxidative phosphorylation, thereby leading to mitochondrial dysfunction and generation of reactive oxygen species (ROS). Given our finding that NR-6226C treatment results in activation of genes involved in oxidative damage response, we tested whether NR-6226C induces formation of ROS in *C. glabrata*. Fungal cells were preloaded with the fluorescent ROS sensor H_2_DCF-DA, after which we measured changes in fluorescence after 3, 6, and 24 hours of exposure to either DMSO, 30 µM NR-6226C, or 75 µM H_2_O_2_ (concentrations that strongly inhibited proliferation; Suppl. Fig. S7A). Surprisingly, even though treatment with NR-6226C for 1h induced the activation of genes involved in the oxidative damage response (Fig. 3A), no ROS formation could be detected after treatment for 4h or 6h (Fig. 4A). Only after 24h of NR-6226C treatment was significant ROS production observed. It is possible that short-term treatment with NR-6226C induces formation of low levels of ROS that are sufficient to induce transcriptional changes but that are below the detection limit of the H_2_DCF-DA assay. It also cannot be excluded that NR-6226C activates the transcriptional oxidative damage response independently of ROS. Nonetheless, these data show that long-term exposure of fungal cells to NR-6226C can induce ROS production.

**Figure 4.**
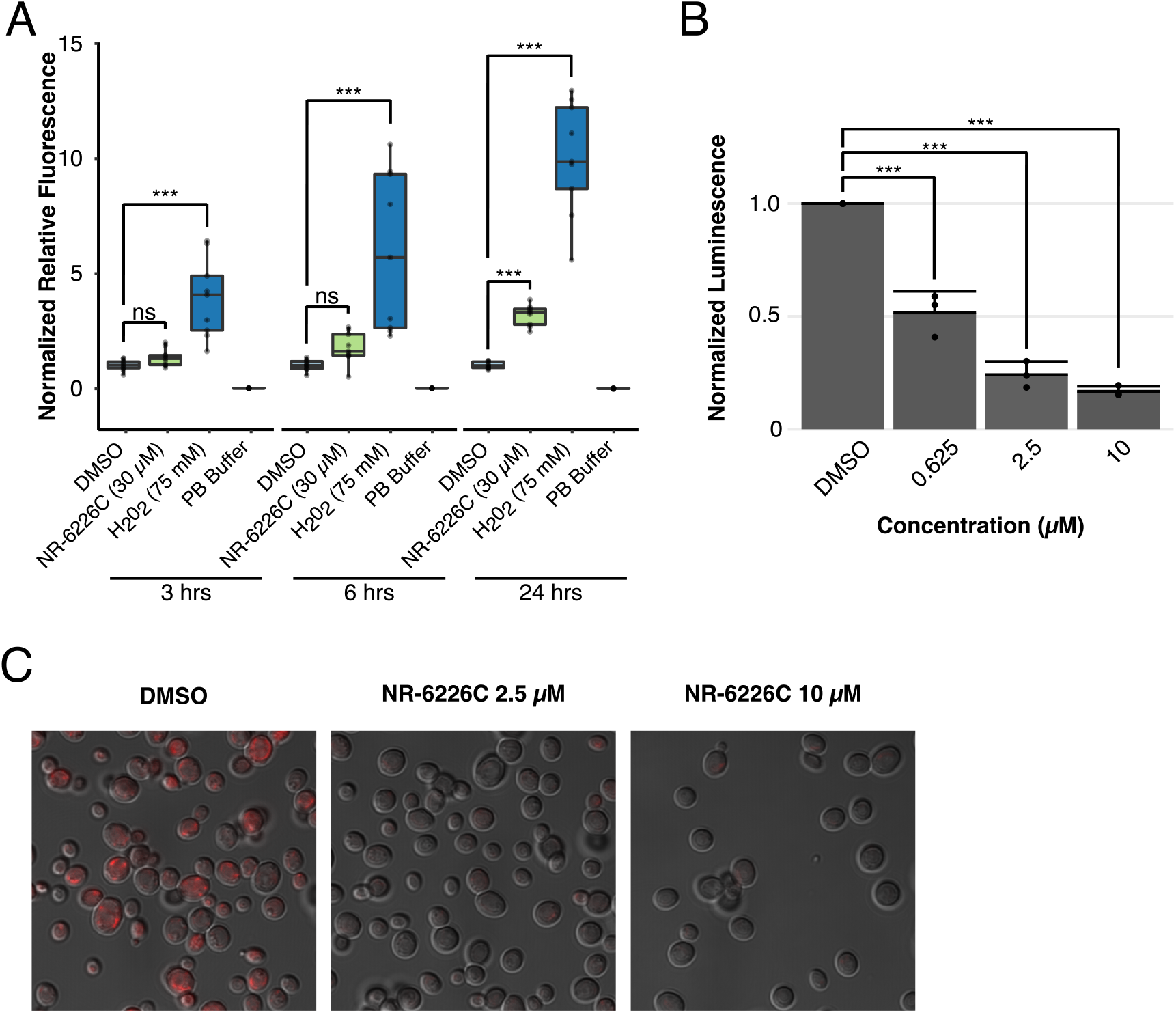
NR-6226C treatment impairs mitochondrial functions and induces ROS formation. ***A,*** *C. glabrata* cells were preloaded with the fluorescent ROS sensor H_2_DCF-DA, after which changes in fluorescence were measured after 3, 6, and 24 hrs after exposure to either DMSO, 30 µM NR-6226C, or 75 µM H_2_O_2_. ***B,*** *C. glabrata* cells were treated for 24 hrs with DMSO, 0.625 µM, 2.5 µM or 10 µM NR-6226C, after which relative ATP levels were measured by luminescence using CellTiter-Glo. An equal number of cells was harvested for each experiment, and luminescence was normalized to DMSO. ***C,*** *C. glabrata* cells were treated with DMSO, 2.5 µM or 10 µM NR-6226C for 24 hrs, after which they were loaded with MitoTracker and imaged using a Zeiss LSM 710 microscope. Representative images of random microscopic fields are shown. \*\*\**P*≤0.001; \*\**P*≤0.01; *ns*, not significant; *n=3*.

We hypothesized that NR-6226C-induced ROS formation was responsible for the antifungal effect of NR-6226C. However, treatment with the ROS scavenger N-acetyl-cysteine (NAC) did not rescue the negative effect of NR-6226C on fungal cell viability (Suppl. Fig. S7B), suggesting that ROS production is not the primary cause of death. Rather, inactivation of crucial iron-dependent metabolic processes may underlie the effect of NR-6226C, such as loss of ATP production and reduced synthesis of various essential metabolites. To test whether NR-6226C affects metabolism and cellular ATP production, we treated *C. glabrata* with increasing concentrations of NR-6226C. Equal numbers of cells were harvested for each concentration point and relative ATP levels were quantified using Cell-Titer Glo^29^. As shown in Figure 4B, treatment with NR-6226C resulted in a concentration-dependent reduction in luminescence, reflecting decreased metabolic activity and loss of cellular ATP production. This suggests that treatment with NR-6226C has a negative effect on mitochondrial activity. We therefore labeled *C. glabrata* cells with the fluorescent dye MitoTracker Red CMXRos, which fluoresces preferentially upon entry into metabolically active mitochondria. Interestingly, after 24h treatment with NR-6226C, cells clearly exhibited lower fluorescence intensities compared to cells treated with DMSO (Fig. 4C), confirming that NR-6226C impairs mitochondrial activity. Taken together, NR-6226C causes significant mitochondrial dysfunction in *C. glabrata* cells, leading to ROS formation, loss of metabolic activity and depletion of cellular ATP levels.

### 26C has antifungal activity in an in vivo infection model

To investigate whether NR-6226C has antifungal activity *in vivo*, we employed *Galleria mellonella* as an infection model. Recently, *G. mellonella* has emerged as a robust infection model for studying virulence and antimicrobial therapies against infectious agents^30^. *Galleria* is capable of mounting an efficient innate immune response against pathogenic fungi that is broadly comparable with the mammalian antifungal immune response^30^. In addition to obvious ethical issues, another important advantage of *G. mellonella* over mice is that there are no concerns of introducing pathogens into breeding facilities, which is a major challenge with mouse models.

In brief, *G. mellonella* larvae were infected with WT *Candida*, followed by incubation in the presence of either DMSO (untreated control) or 30 µM NR-6226C, a concentration well above the *in vitro* IC50 for fungal proliferation. As shown in Figure 5A and B, treatment with NR-6226C of *G. mellonella* larvae infected with either wild-type *C. glabrata* or azole-resistant *C. glabrata* significantly improved the survival of the infected animals. We conclude that NR-6226C is a selective antifungal agent with *in vivo* activity that bypasses azole resistance.

**Figure 5.**
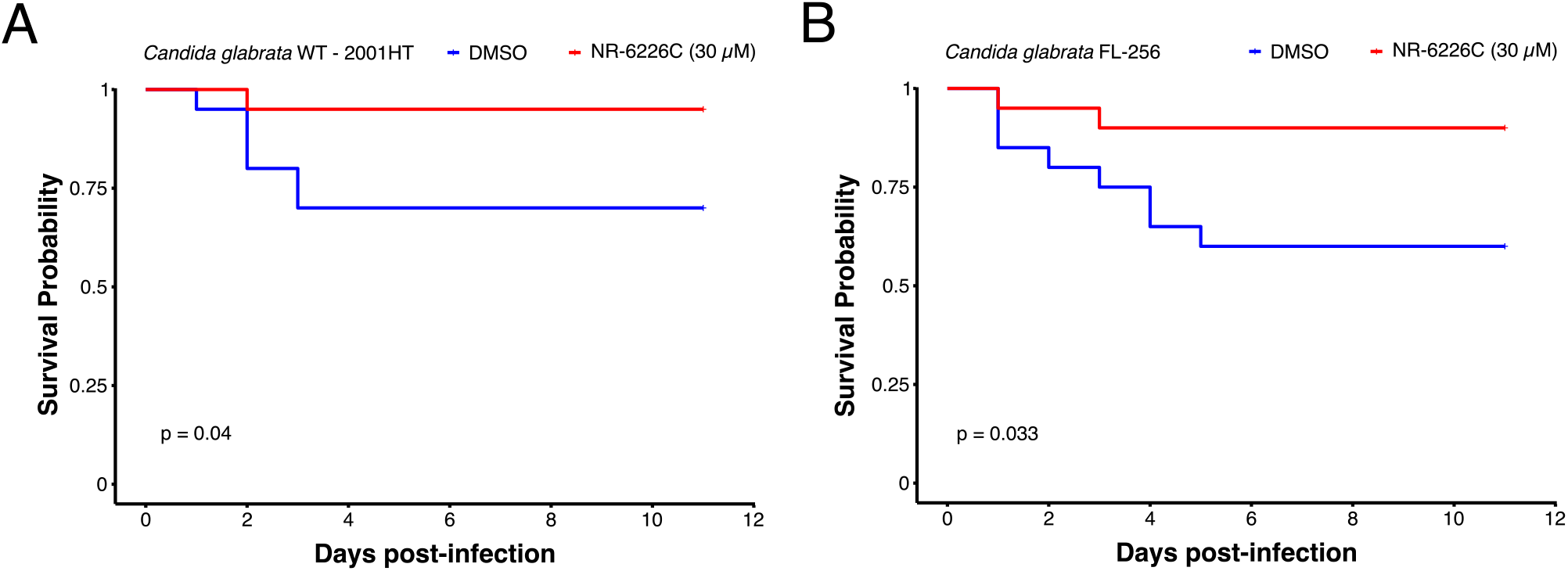
NR-6226C treatment improves the survival of *C. glabrata-infected G. mellonella* larvae. ***A,B,*** *G. mellonella* larvae were infected with either *C. glabrata* 2001HT (A) or *C. glabrata* FL-256 (B), and treated with either DMSO or 30 µM NR-6226C (see Methods), after which overall survival was monitored over time. Survival curves were generated using the Kaplan-Meier formula. Statistical significance was calculated using log-rank tests in R.

### Synergistic antifungal effect of NR-6226C and fluconazole against C. albicans

One approach to countering antifungal resistance that has recently gained interest is combination therapy^31^, where the simultaneous use of two drugs produces either an additive or synergistic effect to impede cellular growth. This strategy could prove beneficial in many circumstances. For example, drugs may be used at lower concentrations to inhibit fungal growth, thereby preventing general toxicity to the host. Combinations of drugs that target different fungal processes may also reduce the likelihood of development of acquired resistance to antifungal drugs. We therefore tested NR-6226C in combination with fluconazole, a first line antifungal that is often clinically ineffective against common nosocomial infections, such as certain *Candida spp*. Interestingly, while wild-type *C. albicans* was only susceptible to high concentrations of fluconazole, combination of fluconazole with NR-6226C resulted in a significantly stronger impairment of fungal proliferation than either drug alone (Fig 6A). To further determine whether the combinatorial effects of these compounds were synergistic, we analyzed our data using the SynergyFinder Plus package^32^ for R using the Bliss model for synergistic interactions^33^. This revealed strong synergy between NR-6226C and fluconazole (Fig. 6B and Supp. Fig. S8A). Although *C. glabrata* is more sensitive to NR-6226C than *C. albicans*, we did not observe substantial additive or synergistic effects of drug combinations (Suppl. Fig. S8B-G). This might be expected, because *C. glabrata* exhibits intrinsically low susceptibility to fluconazole^34^. While analysis of combinations between NR-6226C and a broader panel of antifungal drugs against a spectrum of pathogenic fungi will be the focus of future studies, these results demonstrate that NR-6226C strongly potentiates the antifungal activity of fluconazole against *C. albicans*.

**Figure 6.**
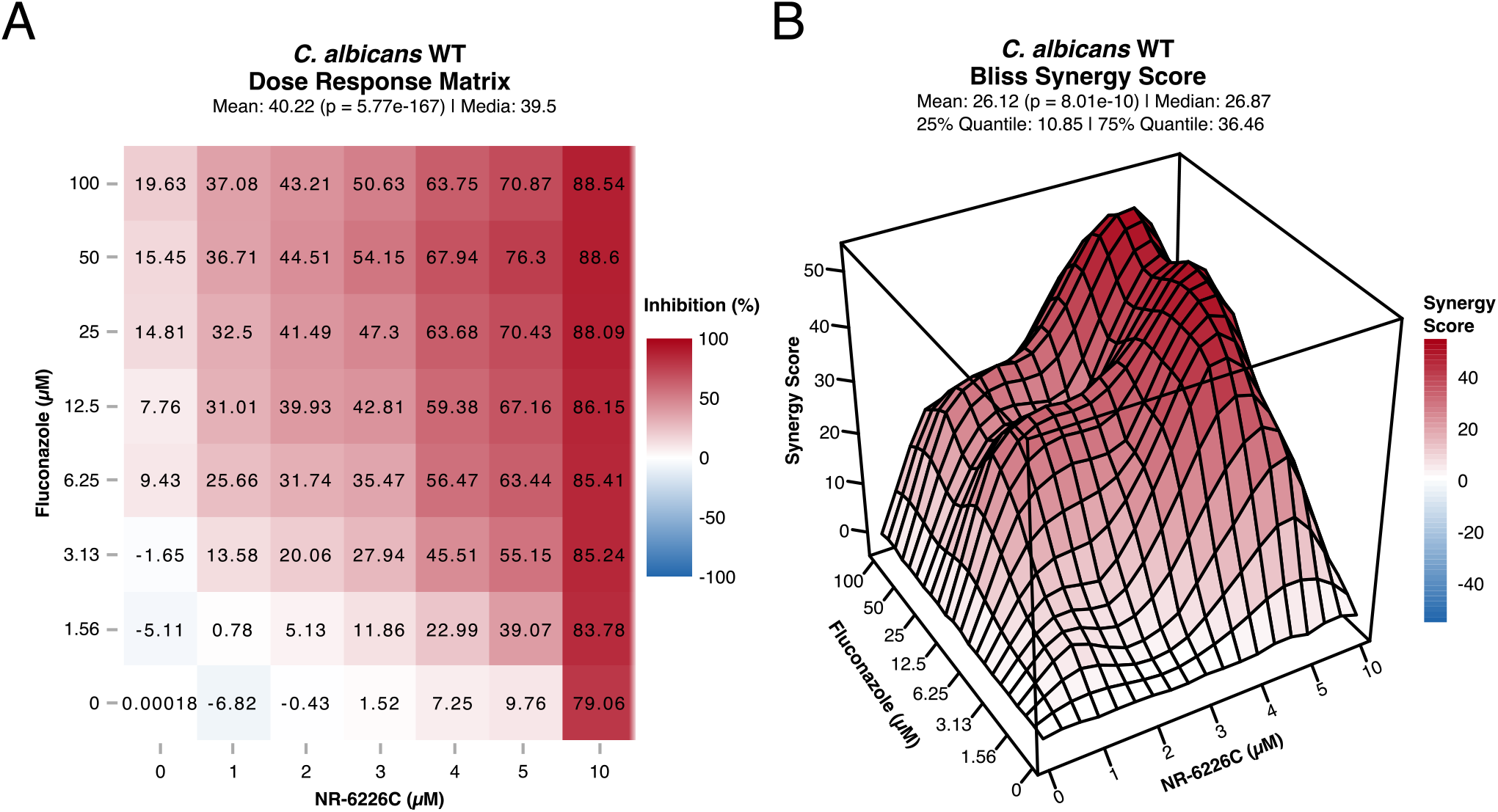
Fluconazole and NR-6226C have a synergistic effect. ***A,*** Dose response matrix of *C. albicans* WT cells treated in combination with Fluconazole and NR-6226C. Relative cell numbers were quantified as described in Figure 1B. ***B,*** 3D visualization of the synergy matrix shown in (A) using the Bliss synergy model. Panel A and B were generated using SynergyFinder in R.

## Discussion

A key challenge in combating fungal resistance is the limited selection of antifungal drugs that are clinically available. Our study revealed that the compound NR-6226C has potential as an antifungal agent. Pathogenic fungi require iron for infection, but the concentration of free iron in mammals is very low (`∼`10^-9^ M) due to sequestration by iron-binding and iron-containing compounds^35^. In our study, we found that NR-6226C prevented *Candida* proliferation *in vitro* and *in vivo* by chelating iron. This resulted in activation of iron starvation response genes such as *FTR1*, *HMX1*, *LSO1* and *FET3* in a Atf1/2-dependent manner. *FET3* encodes a high-affinity iron importer, which mediates iron uptake from the extracellular environment^25^ and it has been proposed that *LSO1* encodes a cytoplasmic protein involved in intracellular iron transport^17^. Furthermore, to overcome the limiting iron availability in their host, pathogenic fungi increase the Aft1-dependent expression of the heme oxygenase Hmx1, which facilitates degradation of heme from the host, thereby releasing iron for subsequent metabolic use^36–38^. Our results indicate that iron chelation by NR-6226C has a direct effect on fungal iron supply, thereby affecting iron metabolism and subsequent intracellular regulation.

Other than the activation of genes regulated by the Aft1 iron-sensory system, our RNA sequencing data revealed the downregulation of enzymatic activities and other genes linked to mitochondrial functions. These findings are consistent with previous yeast studies showing that iron homeostasis is tightly coordinated with metabolic processes^39^. Given that multiple iron-dependent metabolic processes occur in mitochondria, cells likely redirect the limited amount of available iron to essential iron-dependent pathways to maintain homeostasis. The significant decrease of the expression of Fe-S cluster-dependent genes encoding *H_2_S-NADP* oxidoreductase, *Sdh2*, and *Aco1* (Fig. 3A), which have known functions in the mitochondrial respiratory chain^40, 41^, supports this hypothesis. Further evidence that NR-6226C perturbs iron homeostasis is provided by induction of *CAGL0L11990g*, a homolog of *Sc GRX4*. Grx4 is a cytosolic monothiol glutaredoxin that not only functions in concert with Fe-S clusters as an iron sensor, but which is also involved in Fe-S cluster biogenesis in the mitochondria^42, 43^. Although we have not studied the molecular mechanism by which iron depletion results in downregulation of these genes in *C. albicans*, studies in *S. cerevisiae* have revealed that this is at least in part due to Cth2-dependent post-transcriptional degradation and translational-repression of these mRNAs^44, 45^. It is not unlikely that similar mechanisms operate in *C. albicans*.

While NR-6226C has a strong preference for Fe^2+^, the addition of Zn^2+^ and Cu^2+^ to the extracellular medium also countered its antifungal activity, even though these metal ions do not efficiently interact with NR-6226 *in vitro* even at high concentrations. Although we cannot exclude the possibility that the antifungal effect of NR-6226C is mediated by sequestration of trace amounts of Zn^2+^ and Cu^2+^ *in vivo*, we believe it is more likely that these metals induce a stress response that compensates for iron chelation by NR-6226C. Indeed, high concentrations of heavy metals cause mismetallation^14, 46^, which can induce an iron starvation response that results in increased uptake of iron from the environment by microorganisms^15^.

It is well known that persistent iron deprivation leads to increased levels of oxidative stress over time^47^. Similarly, long-term exposure to NR-6226C resulted in a significant increase in ROS production. However, ROS production did not appear to be the direct cause for loss of cell proliferation, as treatment with ROS scavengers did not rescue the effects of NR-6226C. Although long-term NR-6226C-induced ROS production may have a deleterious effect on fungal cells, the immediate effects of NR-6226C may rather be due to impaired metabolic activity and ATP production. These findings mirror previous studies that linked iron metabolism and mitochondrial activity to fungal pathogenicity in *C. albicans*, *C. auris*, and *Aspergillus fumagatus*^48–51^. These links between mitochondria and virulence include lipid biosynthesis, mitochondrial fusion, mitochondrial respiration, and calcium signaling, and indicate that therapies affecting mitochondrial function could prove to be an effective form of antifungal therapy. Indeed, a recent study demonstrated that iron homeostasis and mitochondrial activities may be important targets for treatment of *Candida auris*^50^.

An interesting finding of our study is that NR-6226C strongly synergizes with fluconazole in blocking proliferation of *C. albicans*. Drug combinations can have several advantages, such as the need for lower concentrations of drugs to achieve the desired effect, thereby reducing the risk of off-target toxicity towards host cells. Another advantage is that it is likely more difficult for cells to develop simultaneous resistance against two drugs with separate mechanisms of action. Although more comprehensive drug combination testing against a range of pathogenic fungi will be the focus of future studies, it will be particularly interesting to test combinations of NR-6226C with azoles, because these antifungals exert their effect by blocking ergosterol biosynthesis, and expression of genes required for ergosterol synthesis is strongly dependent on iron availability^52^.

In conclusion, NR-6226C is an interesting lead compound for further development into antifungal therapy. In future studies, it will be of interest to study the effects of NR-6226C on other fungal pathogens such as *Aspergillus fumigatus* and *Cryptococcus neoformans*, and studies in mammalian models will be important to further characterize the *in vivo* potential of NR-6226C, and further studies may be needed to optimize its antifungal activity and ADME profile.

## Methods

### Culture Maintenance

*Candida spp* and *S. cerevisiae* strains used in this study are listed in Suppl. Table S1. Cells were stored at −80°C in YPD (1% Yeast extract, 2% Peptone, and 2% Dextrose) media containing 20% glycerol. For experimental use, cells were streaked out on solid YPD plates (with 2% agar) and incubated at 30°C for 2-3 days until single colonies appeared. The solid plates were stored at 4°C for up to three weeks until new cells were streaked out again.

For experiments, yeast cells were routinely cultured in YPD media and incubated at 30°C overnight. The cultures were refreshed the next day for 2-3 hours in YNB-iron (0.69% Yeast Nitrogen Base without Amino Acids and Iron, and 2% Dextrose) media, unless stated, at the same temperature. After incubation, the optical density (OD_600_) of the cell cultures were measured and adjusted to 0.15.

### Compound Screening

The Collismycin compounds were provided by EntreChem (Spain) at stock solutions of 20 mM in dimethyl sulfoxide (DMSO) and stored at −20°C. For serial dilutions of the compounds in clear 96-well plates, intermediate stock concentrations of 100 µM of compound (100 µl for each technical replicate) were prepared in YNB-iron media and 1:1 serial dilutions were performed using a multi-channel pipette. 50 µl of cell culture (OD_600_ = 0.15) were added to the wells containing the compounds, for a total volume of 150 µl. The final concentrations in the wells (with three technical replicates) were equivalent to 0.5 µM, 1 µM, 2.1 µM, 4.2 µM, 8.4 µM, 16.7 µM, 33.3 µM, and 66.6 µM. The OD_600_ of each well was quantified and the plates were incubated inside plastic bags (to prevent evaporation) at 30°C for 24 hours, after which the final OD_600_ was measured.

### Human Cell Lines

HEK-293 and HS-5 cells were stored at −180°C, in freezing medium (50% DMEM, 40% Fetal bovine serum and 10% DMSO) for long-term storage. For experimental use, the cells were thawed and then cultured using Dulbecco’s Modified Eagle’s Medium (DMEM) containing 10% fetal serum albumin and 1% Penicillin/Streptomycin.

### Cell-Titer Glo Assay

1:1 serial dilutions of the compounds were performed in 96-well plates to obtain a final concentration range (three technical replicates each) equivalent to 0.781 µM, 1.562 µM, 3.125 µM, 6.25 µM, 12.5 µM, 25 µM, 50 µM and 100 µM. Each well had a total volume of 100 µl, consisting of 50 µl compound and 50 µl seeded cells (10000 cells per well). The plates were placed inside plastic bags to prevent evaporation and incubated at 37°C, 5% CO_2_ for 24 hours. 100 µl of Cell Titer-Glo 2.0 (Promega, G9243) was added to each well and then measured for luminescence using a Synergy 2 Gen 5 plate reader from BioTek.

### Nano liquid chromatography-mass spectrometry

Liquid chromatography-mass spectrometry (LC-MS) grade water, acetonitrile, and formic acid (≥99%) were purchased from VWR (Radnor, PA, U.S). NR-6226C (Terpyridine (2,2′:6′,2′′-Terpyridine, 98%)) was purchased from Sigma Aldrich (now Merck, KGaA, Darmstadt Germany). Metal solutions containing 2 µM, 4 µM, 6 µM and 10 µM CaCl_2_, CuCl_2_, FeCl_2_, FeCl_3_, MgCl_2_, MnCl_2_, and ZnCl_2_ in 0.1 % formic acid in LC-MS grade water (hereby referred to as only 0.1 % formic acid) was made. A stock solution of terpyridine (100 µM) in 0.1% formic acid was made and added to the metal solutions to a final concentration of 10 µM terpyridine. All metal solutions in addition to a blank sample containing 10 µM terpyridine in 0.1 % formic acid were subjected to nano liquid chromatography (nanoLC-MS) analysis. NanoLC-MS was performed using an nLC EASY 1000 pump connected to a Q-Exactive mass spectrometer (MS) with a Nanospray Flex^TM^ ion source (all from Thermo Fisher Scientific, Waltham, MA, U.S). For separation, a 50 µm (inner diameter) x 5 cm in-house packed nanoLC column containing 2.6 µm Accucore C18 particles (80 Å) was used. The column was packed according to the protocol from Berg, et.al^56^. Trapping of analytes prior to separation was performed on-line using a 2 cm Acclaim PepMap with 3 µm particles (100 Å). All columns and packing materials were obtained from Thermo Fisher Scientific. The mobile phases consisted of 0.1% formic acid (A) and 90/10/0.1 acetonitrile/water/FA (v/v/v) (B). A linear gradient from 5-20% B in 10 min was employed, and trapping was performed by 100% mobile phase A. Equilibration was performed using 2 µl and 3 µl 100% A for the trapping column and analytical column, respectively. The flow rate was set to 250 nl/min and the injection volume used was 1 µl.

The MS was operated in positive mode using single ion monitoring (SIM). A mass-to-charge ratio (*m/z*) of 234.1 (M+H) was used for terpyridine and an *m/z* of 261.1 (2M+Fe)^2+^ was used for terpyridine bound to Fe^2+^. All data processing was performed by using the XCalibur^TM^ Software (Thermo Fisher Scientific).

### Cell proliferation with metal supplementation

NR-6226C (Sigma-Aldrich) was prepared at a stock solution of 20 mM. 1 mM stocks of metal solutions (CaCl_2_, CuCl_2_, FeCl_2_, FeCl_3_, MgCl_2_, MnCl_2_, and ZnCl_2_) were prepared in mQH_2_O and autoclaved. Serial dilutions of NR-6226C using YNB-iron media, were performed in clear 96-well plates to obtain a final concentration range of 0.78 µM, 1.56 µM, 3.13 µM, 6.25 µM, 12.5 µM, 25 µM, 50 µM and 100 µM. 50 µl of either mQH_2_O or metal solution were added to each well, including 50 µl of refreshed *C. glabrata* WT with an adjusted OD_600_ of 0.15. The 96-well plates were measured at OD_600_ and incubated in plastic bags to prevent evaporation for 24 hours, after which the final OD_600_ was measured.

### RNA Sequencing

Cells from overnight precultures in YPD medium were reinoculated in fresh YNB medium without iron and grown until OD_600_ = 0.4. Cultures were then treated for 1 hr with DMSO (vehicle) or 10 µM NR-6226C or 5 µM Dp44mT. Cells were collected from three independent experiments, snap frozen and stored at −80°C. Total RNA purification was performed as previously described using the RNeasy Mini Kit (74104, Qiagen) and gDNA removal was made using the RNase-Free DNase Set kit (79254, QIAGEN). RNA quantification and quality control were done with a Tape Station 4150 (Agilent)^57^.

Library preparation and sequencing were performed at GenomEast (IGBMC, Illkirch, France). One DMSO 0.05% sample was excluded from the analysis due to insufficient quality. The library was sequenced on Illumina Hiseq 4000 sequencer as Single-Read 50 base reads following Illumina’s instructions. Reads were pre-processed in order to remove adapter, polyA and low-quality sequences (Phred quality score below 20). Reads shorter than 40 bases were discarded from further analysis. These pre-processing steps were performed using cutadapt version 1.10^58^. Image analysis and base calling were performed using RTA 2.7.7 and bcl2fastq2.17.1.14. Adapter dimer reads were removed using DimerRemover (https://sourceforge.net/projects/dimerremover/). The quality of the RNAseq reads was examined using the FastQC 0.11.2 (http://www.bioinformatics.babraham.ac.uk/projects/fastqc/) and FastQScreen 0.5.1 (http://www.bioinformatics.babraham.ac.uk/projects/fastq screen/). Reads were mapped onto the ASM254v2 assembly of *Candida glabrata* genome using STAR version 2.5.3a^59^. Gene expression quantification was performed from uniquely aligned reads using htseq-count version 0.6.1p1^60^, with annotations from Ensembl fungi version 50 and union mode. Only non-ambiguously assigned reads have been retained for further analyses. Read counts were normalized across samples with the median-of-ratios method^61^. Comparisons were implemented in the Bioconductor package DESeq2 version 1.16.1 using the test for differential expression^62^. Genes with no p-value corresponded to genes with high Cook’s distance that were filtered out. P-values were adjusted for multiple testing using the Benjamini and Hochberg method^63^. Genes with no adjusted p-value correspond to genes filtered out in the independent filtering step in order to remove reads from genes that have no or little chance of showing significance evidence of differential expression.

### RT-qPCR

WT and *atf1*Δ cells were grown overnight in CSM. Cells were then diluted 10 times and grown until log phase at which point 10 µM of NR6226C or Dp44mt were added to the medium, either in absence or presence of 5 µM of FeCl_2_ as indicated. After 24 hrs of incubation cells were collected and total RNA purification, reverse transcription and RT-qPCR were performed as previously described with minor modifications^57, 64^. Briefly, total RNA was purified using the RNeasy Mini Kit (74104, Qiagen) and reverse transcription was performed using the QuantiTect Reverse Transcription Kit (205311, Qiagen). RT-qPCR experiments were done using the HOT FIREPol® EvaGreen® qPCR Mix (08-36-00001-10, Solis Biodyne) and a LightCycler® 96 System (Roche). Expression of the 5S gene was used as a control. Primers used in RT-qPCR experiments:

*FTR1*: GATTGGGTTCTTGAGTAGAAG and GAGCCCTGTGTGGTAATA

*FET3*: GTCAATATGAAGACGGGATG and CCACTCACTAAGCGATAAAG

*HMX1*: CCAGAGATGCCCACAATA and CATAATAGTACGCCAGAATACC

*LSO1*: AGAAGAAGCTGATTATGGAAC and CTGCCTCCCTTACCTAAA

*5S*: GTTGCGGCCATATCTACCAGAAAG and CGTATGGTCACCCACTACACTACT

### Reactive oxygen species (ROS) assay

Our method was adapted using the protocol developed by James, et al.^65^ for ROS assessment in yeast. The overnight culture was refreshed in YNB-iron media for 2-3 hours and then adjusted to OD_600_ = 0.5 in 100 ml. The cells were washed by centrifugation for 5 minutes at 4300 rpm, followed by aspiration of the supernatant and a re-suspension of the cell pellets in 20 ml phosphate buffer solution (PB, 0.1M, pH 7.4). 1 ml of cell suspension was aliquoted to Falcon tubes and then centrifuged again to remove the supernatant. The general oxidative stress indicator, CM-H_2_DCFDA (Thermo Fisher), was prepared in a stock solution of 1 mM DMSO and then diluted to 10 µM using PB solution. The cell pellets were pre-loaded with the dye by re-suspension in 10 µM CM-H_2_DCFDA and then incubated in the dark at 30°C for 30 minutes. Cell pellets that were used for a negative control of the dye were treated with PB solution alone. After incubation, the samples were centrifuged at 4300 rpm for 2 minutes and followed by aspiration of the supernatant. The cell pellets were then treated with either DMSO, 30 µM NR-6226C, or 75 mM hydrogen peroxide (H_2_O_2_), including the negative controls. The samples were placed in a 30°C incubator and then measured for OD_600_ after 3, 6 and 24 hours of incubation, after which an equal number of cells (OD_600_ = 0.5, 1 ml) were harvested. The harvested cells were centrifuged at 6000g for 10 minutes, with the supernatant aspirated and the cells resuspended in PB solution. The fluorescence of the samples was measured in a black, clear 96-well plate using a spectrofluorometer.

### Cell-Titer Glo assay in *Candida*

A volume of 500 ml cell culture (OD_600_ = 0.15) was mixed to a final concentration of 1 ml DMSO, 0.625 µM NR-6226C, 2.5 µM NR-6226C, or 10 µM NR-6226C. The samples were incubated for 24 hours at 30°C and then measured for OD_600_. Equal number of cells were harvested in each experiment according to an OD_600_ = 0.1-0.4, then centrifuged at 7000 rpm for 5 minutes to remove the supernatant. The cell pellets were resuspended in 100 µl and then transferred to a 96-well plate. Finally, the samples were treated with 100 µl Cell-Titer Glo and then measured for luminescence using a plate reader.

### Mitochondrial staining with MitoTracker

MitoTracker Red CMXROS (Invitrogen) was prepared to a stock solution of 1 mM using DMSO and stored at −20°C. 5 ml *C. glabrata* WT cells of OD_600_ = 0.15 were centrifuged at 7000 rpm for 5 minutes to obtain cell pellets. The cell pellets were resuspended in 5 ml DMSO, 2.5 µM or 10 µM NR-6226C and then incubated at 30°C for 24 hours. The samples were measured for OD_600_, followed by harvesting of cells to an OD_600_ = 0.3 and then centrifuged at 7000 rpm for 5 minutes.

The cell pellets were resuspended in 100 nM MitoTracker and incubated at 37°C for 25 minutes. The samples were centrifuged at 7000 rpm for 5 minutes and the resulting pellets were washed with 1 ml milliQ H_2_O. The cell pellets were resuspended in milliQ H_2_O and then imaged using a Zeiss LSM 710 microscope.

### Survival assays in infected *Galleria mellonella* models

*Galleria mellonella* was obtained from R.J. Mous Livebait (The Netherlands). The larvae were selected by weight (0.2-0.3 grams) and by the absence of dark spots on the cuticle. The larvae were maintained at room temperature, and the day before the experiment they were transferred to the temperature at which the experiment was going to be performed (30° C). The number of dead caterpillars was scored every day. A group of 20 larvae were incubated with PBS, DMSO and with NR-6226C alone as controls in every experiment. *G. mellonella* were infected with either *C. glabrata* 2001HT or FL-256; *C. glabrata* cells were grown in liquid media (Oxoid) overnight at 30° C with moderate shaking (150 rpm). Groups of 20 larvae per strain were infected in each experiment. The pro-leg area was cleaned with 70% ethanol using a swab. The larvae were inoculated with 10 µl of *C. glabrata* suspension at 2.5 x 10^6^ cells/ml in PBS containing 50 µg/ml of ampicillin by injection, using a 26-gauge needle with Hamilton syringes in the last left proleg. Within 2 hours of infection, 10 µl of compound solution NR-6226C (15 and 30 µM) was injected into a different pro-leg using the same technique.

### Combination treatment with Fluconazole and NR-6226C

Fluconazole (TargetMol) and NR-6226C were prepared to a stock solution of 20 mM using DMSO. The drug and compound were dispensed into clear 96-well plates using CERTUS FLEX liquid dispenser at a concentration range of 1.56 µM, 3.12 µM, 6.25 µM, 12.5 µM, 25 µM, 50 µM and 100 µM for Fluconazole and 1 µM, 2 µM, 3 µM, 4 µM, 5 µM and 10 µM for NR-6226C. 50 µl of refreshed *Candida* cells equivalent to OD_600_ = 0.15 were added to the wells and measured for OD_600_. The plates were placed in plastic bags and then incubated at 30°C for 24 hours until the final OD_600_ was measured again.

## Data Availability

Data description: RNAseq dataset. Name of the repository: ENA database. Accession number: PRJEB63373.

## Acknowledgments

This work was supported by grants from the Norwegian Cancer Society (project numbers 182524 and 208012), the Norwegian Health Authority South-East (2017064, 2017072, 2018012, 2019096), the Research Council of Norway (261936, 301268, 262652). OZ is supported by grant PID2020-114546RB by MCIN/AEI/10.13039/501100011033. Sequencing was performed by the GenomEast platform, a member of the ‘France Genomique’ consortium [ANR-10-INBS-0009]. We thank Cecilie Torp Andersen (Oslo University Hospital) for providing patient-derived *Candida* strains that were used in this work. We also thank all members of the Enserink group and the Knævelsrud group for helpful discussions and feedback.

## Supplemental Figure Legends

**Supplemental Figure S1.** Collismycin-related compound names and structures.

**Supplemental Figure S2.** Compound screening. Heatmap of relative *Candida* proliferation after treatment for 24 hrs with the indicated concentrations of Collismycin analogs. Cell growth was measured using OD_600_. Data were normalized to cells treated with DMSO.

**Supplemental Figure S3. *A, B,*** Proliferation curves and EC_50_ values of *Candida spp* treated for 24 hrs with either Collismycin A (A) or with NR-6226C (26C), NR-6226K (26K), NR-6226V (26V) (B). Cell growth was measured using OD_600_. ***C,*** Proliferation curves and EC_50_ values of HEK-293 and HS-5 cells after treatment for 24 hrs with either NR-6226C, NR-6226K, or NR-6226V. Closed circles indicate technical replicates, lines indicate biological replicates. EC_50_ values were calculated as described in Figure 2C.

**Supplemental Figure S4.** HEK-293 and HS-5 cells were treated with either DMSO, 100 µM NR-6226C, or 100 µM NR-6226K and imaged using the Incucyte Live-Cell Analysis System. Red arrows indicate compound precipitation.

**Supplemental Figure S5.** Mass spectroscopy of NR-6226C on its own and or bound to Fe^2+^.

**Supplemental Figure S6. *A,*** Volcano plot showing commonly regulated genes based on the threshold for the calculated mean difference and standard deviation in log fold-change between *C. glabrata* cells treated with either NR-6226C or Dp44mT (log2-fold change <-0.683 or >0.635). ***B,C,*** Venn diagram showing the count and respective percentages of up-regulated (B) or down-regulated (C) genes between *C. glabrata* cells treated with either NR-6226C or Dp44mT.

**Supplemental Figure S7. *A,*** Growth curves of *C. glabrata* cells pre-loaded with the H_2_DCFDA and then treated with DMSO, 30 µM NR-6226C, or 75 µM H_2_O_2_. Cell proliferation was measured and corrected using OD_600_. ***B,*** EC_50_ values of *C. glabrata* cells treated with NR-6226C alone within a concentration range of 0.78µM-100 µM, or with NR-6226C in combination with 1 µM, 5 µM or 10 µM N-Acetyl-L-Cysteine (NAC). Samples were normalized to DMSO controls, followed by EC_50_ estimation. Statistical significance was calculated using unpaired two-sample t-tests. \*\*\**P*≤0.001; \*\**P*≤0.01; *ns*, not significant; *n=3*.

**Supplemental Figure S8. *A, C, E, G,*** Bliss synergy scores of *Candida spp.* treatment with Fluconazole and NR-6226C combination. ***B, D, F,*** Dose response matrix of *Candida glabrata* cells treated in combination with Fluconazole and NR-6226C. Relative cell numbers were quantified as described in Figure 1B. Dose response matrices and synergy scores were obtained using SynergyFinder in R.

