## Supplemental Figure S1 for "Characterization of a selective, iron-chelating antifungal compound that disrupts fungal metabolism and synergizes with fluconazole"

### Supplemental Figure 1

| Compound | Structure |
|---|---|
| Collismycin 21 | 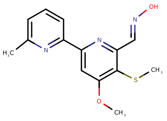 |
| Collismycin 22 | 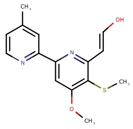 |
| Collismycin 22-ACID | 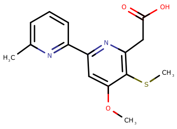 |
| Collismycin A | 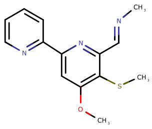 |
| Collismycin DC | 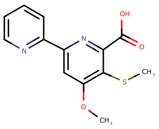 |
| Collismycin DH | 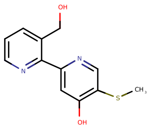 |
| Collismycin H | 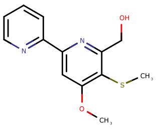 |
| Collismycin H-BUT | 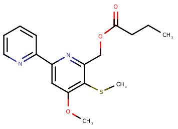 |
| Collismycin HA | 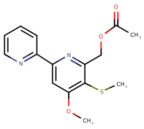 |
| Collismycin SN | 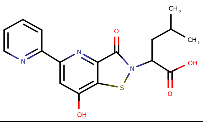 |
| Collismycin SC | 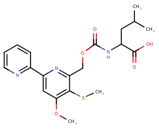 |
| NR-4492C | 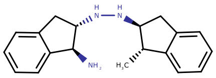 |
| NR-4493C | 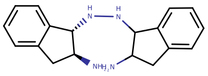 |

| Compound | Structure |
|---|---|
| NR-5012 | 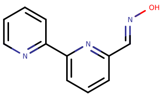 |
| NR-6226A | 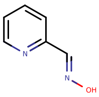 |
| NR-6226B | 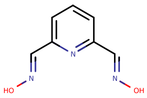 |
| NR-6226C | 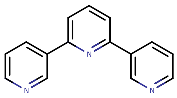 |
| NR-6226D | 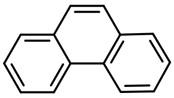 |
| NR-6226K | 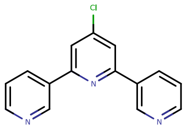 |
| NR-6226V | 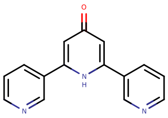 |
| NR-6266A | 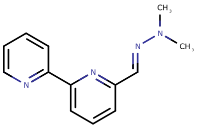 |
| NR-6266B | 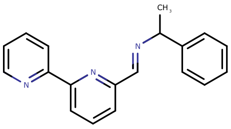 |
| NR-6265P | 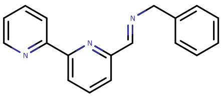 |
| NR-6268A | 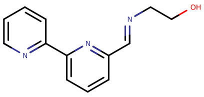 |
| NR-6269A | 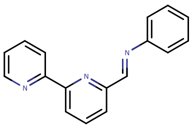 |
| NR-6269B | 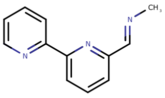 |

**Supplemental Figure S1.** Collismycin-related compound names and structures.
