## Supplemental Figure S2 for "Characterization of a selective, iron-chelating antifungal compound that disrupts fungal metabolism and synergizes with fluconazole"

### Supplemental Figure 2

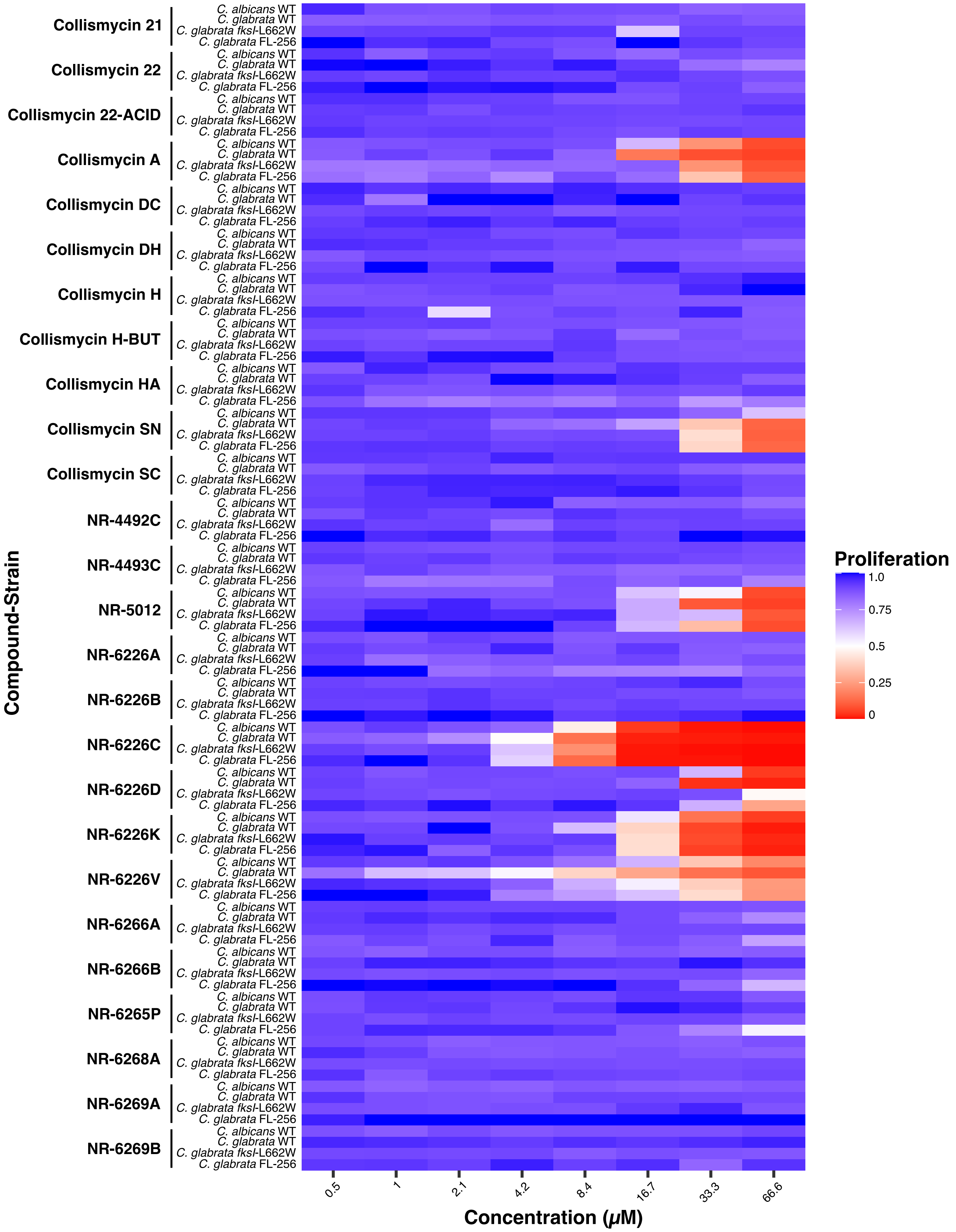


**Supplemental Figure S2.** Compound screening. Heatmap of *Candida* proliferation after treatment for 24 hrs with the indicated concentrations of Collismycin analogs. Cell growth was measured using $OD_{600}$. Data were normalized to cells treated with DMSO.
