## Supplemental Figure S3 for "Characterization of a selective, iron-chelating antifungal compound that disrupts fungal metabolism and synergizes with fluconazole"

### Supplemental Figure 3

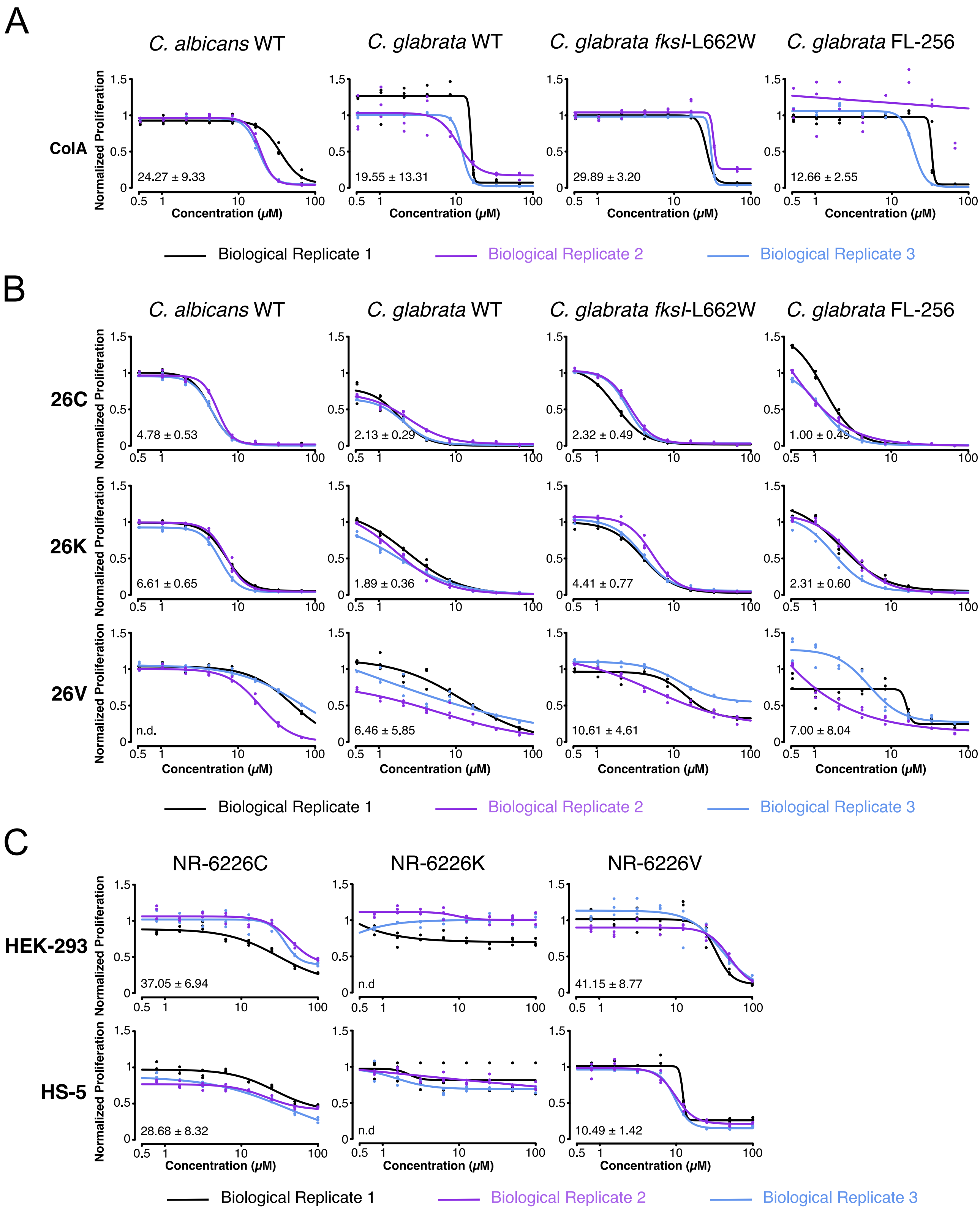


**Supplemental Figure S3.** ***A, B,*** Proliferation curves and $EC_{50}$ values of *Candida spp* treated for 24 hrs with either ColA (A) or with NR-6226C (26C), NR-6226K (26K), NR-6226V (26V) (B). Cell growth was measured using $OD_{600}$. ***C,*** Proliferation curves and $EC_{50}$ values of HEK-293 and HS-5 cells after treatment for 24 hrs with either NR-6226C, NR-6226K, or NR-6226V. Closed circles indicate technical replicates, lines indicate biological replicates. $EC_{50}$ values were calculated as described in Figure 2C.
