## Supplemental Figure S4 for "Characterization of a selective, iron-chelating antifungal compound that disrupts fungal metabolism and synergizes with fluconazole"

### Supplemental Figure 4

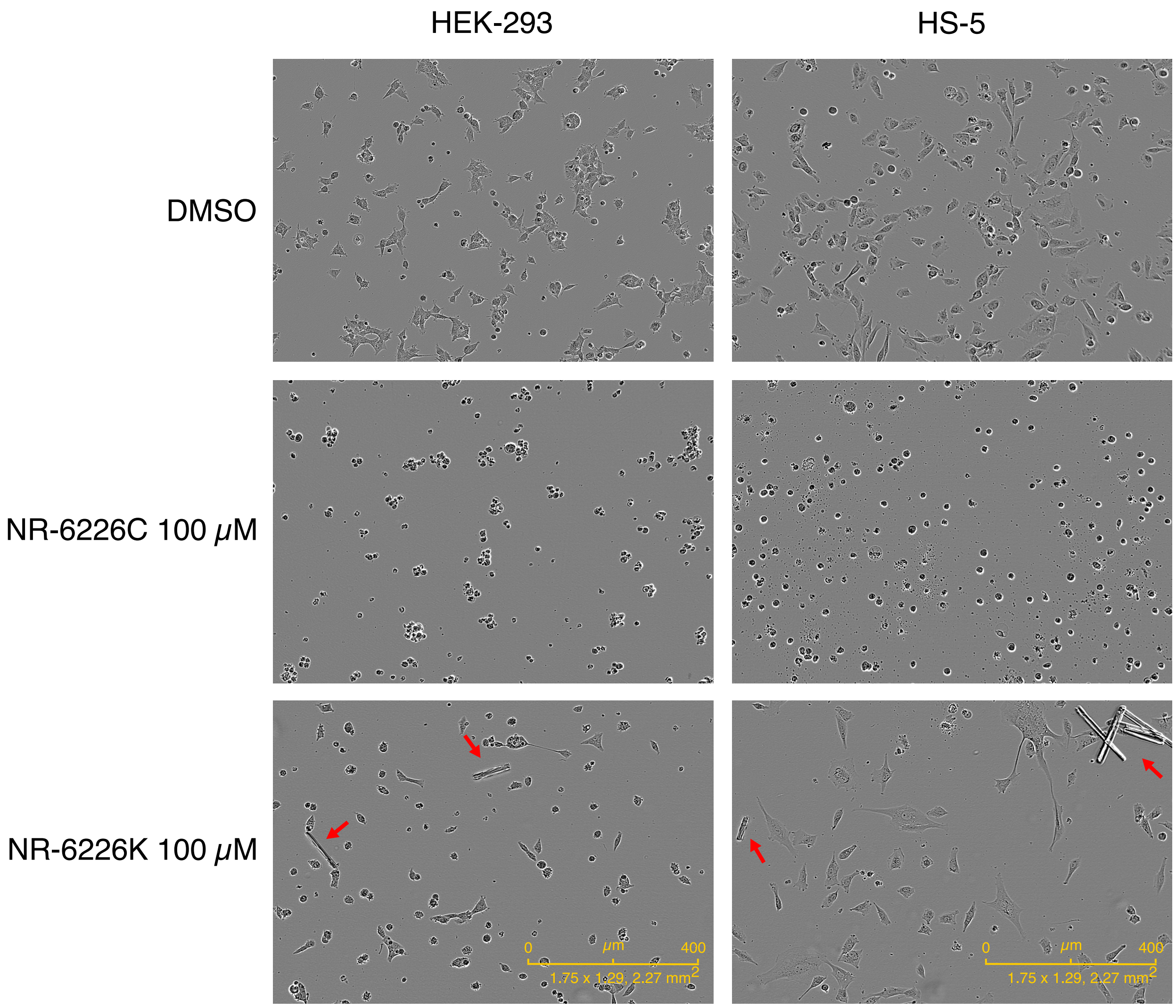


**Supplemental Figure S4.** HEK-293 and HS-5 cells were treated with either DMSO, 100 μM NR-6226C, or 100 μM NR-6226K and imaged using the Incucyte Live-Cell Analysis System. Red arrows indicate compound precipitation.
