## Supplementary figures and images for "Characterization of a selective, iron-chelating antifungal compound that disrupts fungal metabolism and synergizes with fluconazole"

### Supplemental Figure S5

#### Supplemental Figure 5

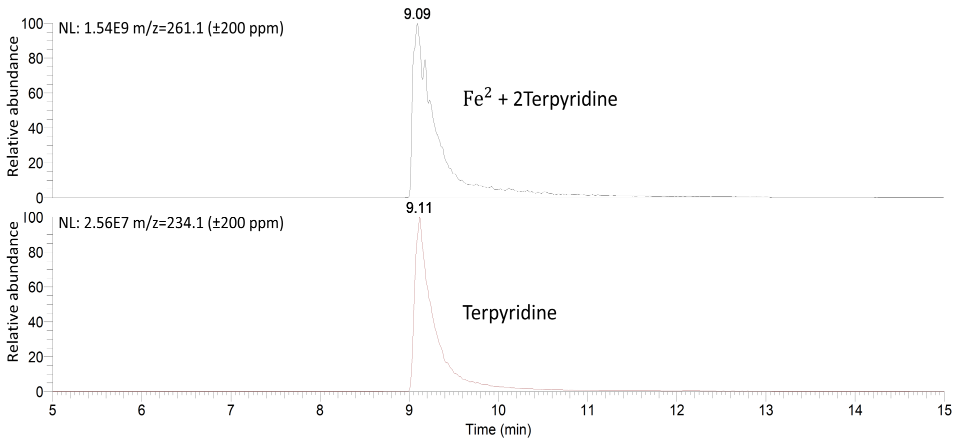


**Supplemental Figure S5.** Mass spectroscopy of 26C on its own and or bound to $Fe^{2+}$.
