## Supplemental Figure S6 for "Characterization of a selective, iron-chelating antifungal compound that disrupts fungal metabolism and synergizes with fluconazole"

### Supplemental Figure 6

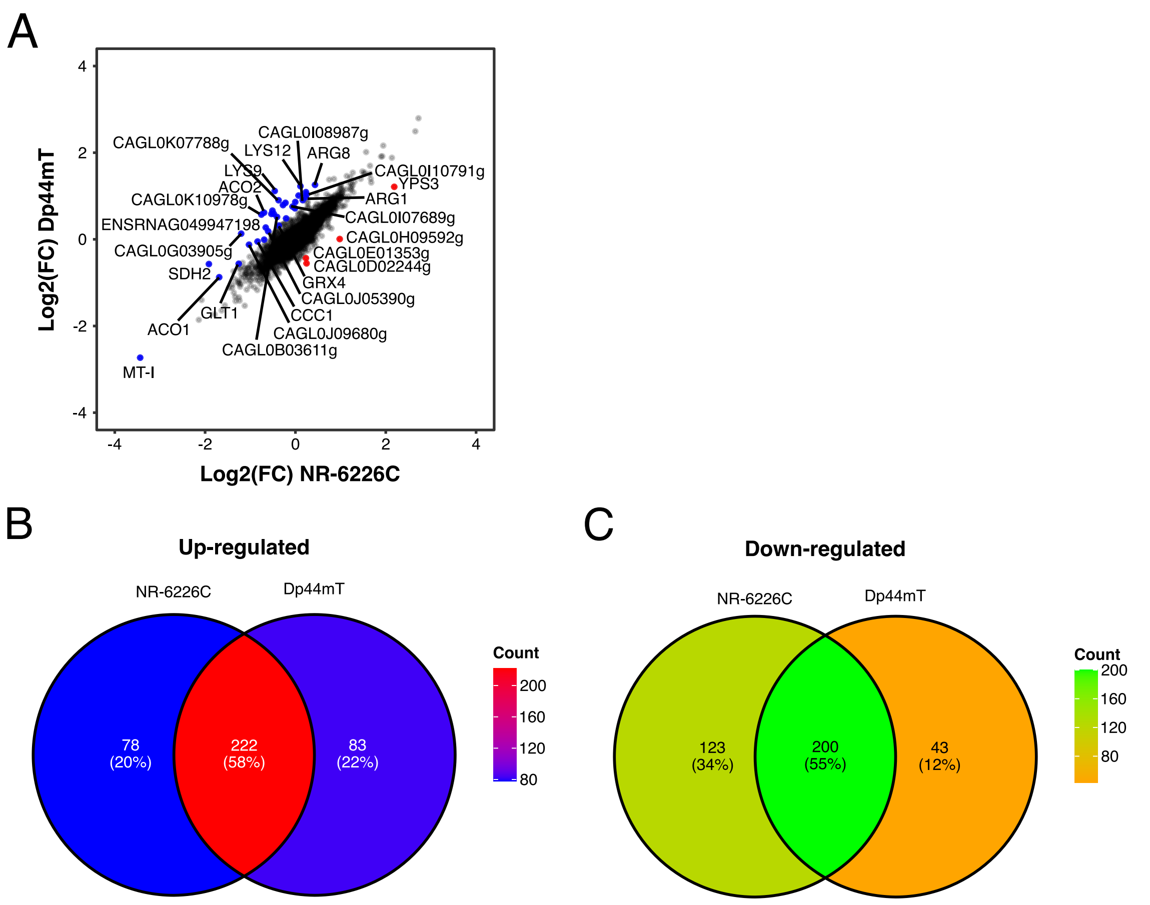


**Supplemental Figure S6.** ***A,*** Volcano plot showing commonly regulated genes based on the threshold for the calculated mean difference and standard deviation in log fold-change between *C. glabrata* cells treated with either NR-6226C or Dp44mT (log2-fold change <-0.683 or >0.635). ***B,C,*** Venn diagram showing the count and respective percentages of up-regulated (B) or down-regulated (C) genes between *C. glabrata* cells treated with either NR-6226C or Dp44mT.
