## Supplemental Figure S7 for "Characterization of a selective, iron-chelating antifungal compound that disrupts fungal metabolism and synergizes with fluconazole"

### Supplemental Figure 7

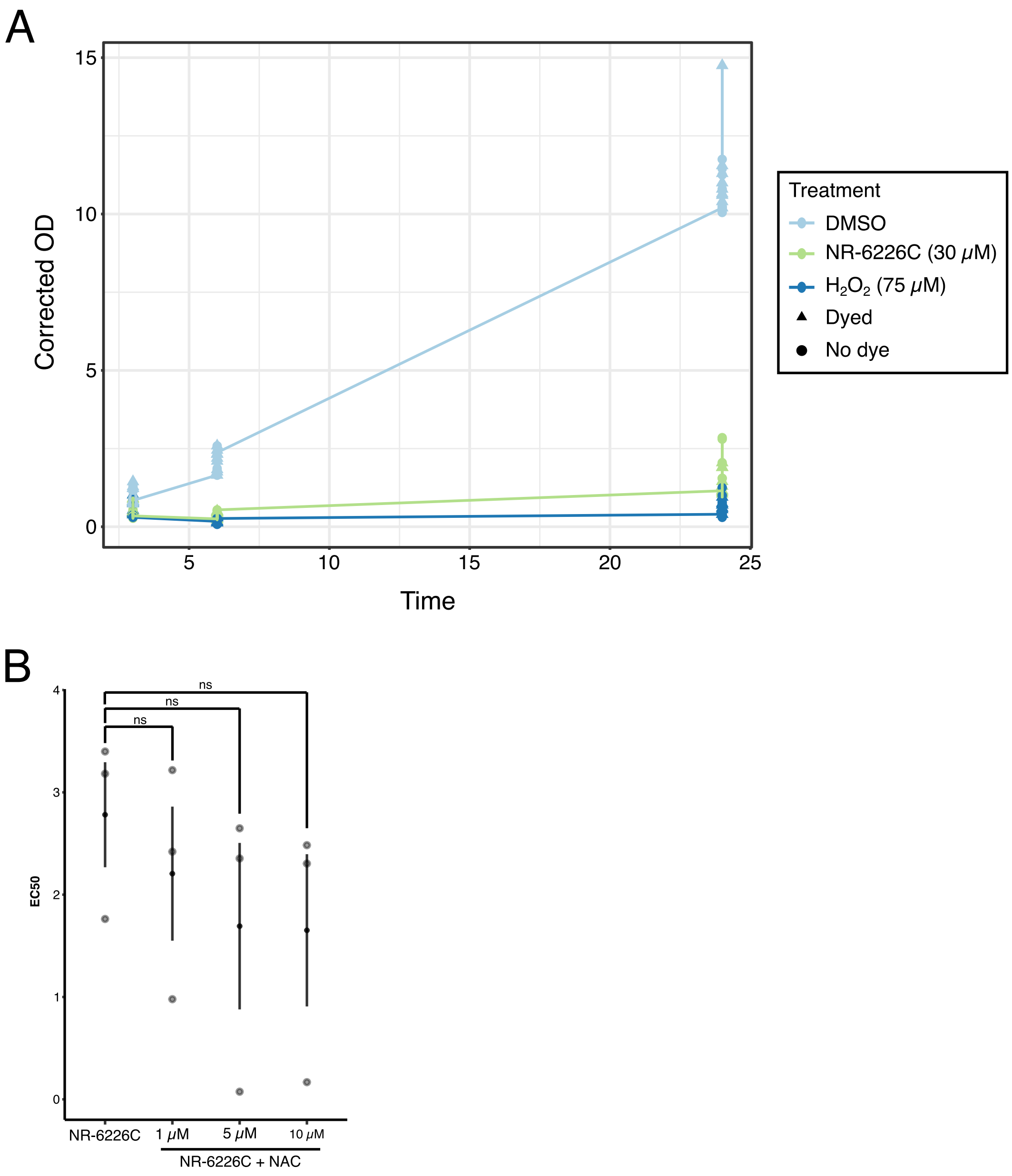


**Supplemental Figure S7.** ***A,*** Growth curves of *C. glabrata* cells pre-loaded with the $H_2$DCFDA and then treated with DMSO, 30 $\mu$M NR-6226C, or 75 $\mu$M $H_2O_2$. Cell proliferation was measured and corrected using $OD_{600}$. B, $EC_{50}$ values of *C. glabrata* cells treated with 26C alone within a concentration range of 0.78$\mu$M-100 $\mu$M, or with NR-6226C in combination with 1 $\mu$M, 5 $\mu$M or 10 $\mu$M N-Acetyl-L-Cysteine (NAC). Samples were normalized to DMSO controls, followed by $EC_{50}$ estimation. Statistical significance was calculated using unpaired two-sample t-tests. ****P≤0.001; **P≤0.01; ns, not significant; n=3.*
