## Supplemental Figure S8 for "Characterization of a selective, iron-chelating antifungal compound that disrupts fungal metabolism and synergizes with fluconazole"

### Supplemental Figure 8

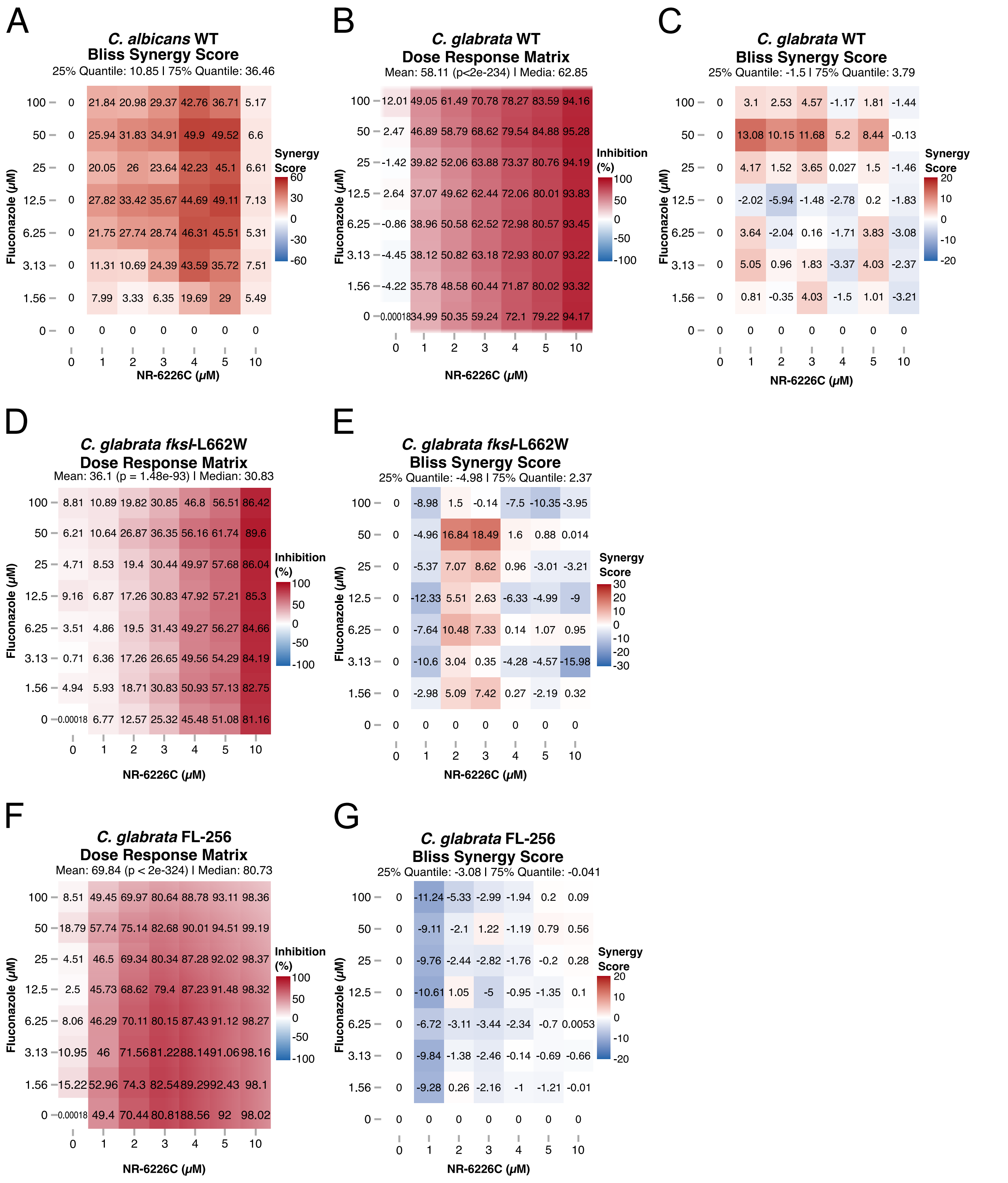


**Supplemental Figure S8.** ***A, C, E, G,*** Bliss synergy scores of *Candida spp.* treatment with Fluconazole and NR-6226C combination. ***B, D, F,*** Dose response matrix of *Candida glabrata* cells treated in combination with Fluconazole and NR-6226C. Relative cell numbers were quantified as described in Figure 1B. Dose response matrices and synergy scores were obtained using SynergyFinder in R.
