## Supplemental Table S1 for "Characterization of a selective, iron-chelating antifungal compound that disrupts fungal metabolism and synergizes with fluconazole"

**Supplemental Table S1.** Yeast strains.

| Species | Strain number | Relevant characteristics or genotype | Source |
|---|---|---|---|
| *C. albicans* | JEY13033 | N/A | CCUG32723 |
| *C. glabrata* | JEY10028 | N/A | ATCC15545 |
| *C. glabrata* | JEY12725 | *fks1*-L662W; Patient-derived from Oslo University Hospital | This study |
| *C. glabrata* | JEY12726 | FL-256; Patient-derived from Oslo University Hospital | This study |
| *C. glabrata* | JEY12527 | 2001HT; *his3Δ trp1Δ* | Kitada, et al. (1995)[53] |
| *S. cerevisiae* | BY4741 | *MATa his3Δ1, leu2Δ0, met15Δ0, ura3Δ0* | Brachmann, et al. (1998)[54] |
| *S. cerevisiae* | *aft1*Δ | *MATa his3Δ1, leu2Δ0, met15Δ0, ura3Δ0, aft1::KanMX* | Giaever, et al. (2002)[55] |
